# High diversity and variability of pipolins among a wide range of pathogenic *Escherichia coli* strains

**DOI:** 10.1101/2020.04.24.059261

**Authors:** Saskia-Camille Flament-Simon, María de Toro, Liubov Chuprikova, Miguel Blanco, Juan Moreno-González, Margarita Salas, Jorge Blanco, Modesto Redrejo-Rodríguez

## Abstract

Self-synthesizing transposons are integrative mobile genetic elements (MGEs) that encode their own B-family DNA polymerase (PolB). Discovered a few years ago, they are proposed as key players in the evolution of several groups of DNA viruses and virus-host interaction machinery. Pipolins are the most recent addition to the group, are integrated in the genomes of bacteria from diverse phyla and also present as circular plasmids in mitochondria. Remarkably, pipolins-encoded PolBs are proficient DNA polymerases endowed with DNA priming capacity, hence the name, primer-independent PolB (piPolB).

We have now surveyed the presence of pipolins in a collection of 2238 human and animal pathogenic *Escherichia coli* strains and found that, although detected in only 25 new isolates (1.1%), they are present in *E. coli* strains from a wide variety of pathotypes, serotypes, phylogenetic groups and sequence types. Overall, the pangenome of strains carrying pipolins is highly diverse, despite the fact that a considerable number of strains belongs to only three clonal complexes (CC10, CC23 and CC32). Comparative analysis with a set of 67 additional pipolin-harboring strains from GenBank further confirmed these results. The genetic structure of pipolins shows great flexibility and variability, with the piPolB gene and the attachment sites being the only common features. Most pipolins contain one or more recombinases that would be involved in excision/integration of the element in the same conserved tRNA gene. This mobilization mechanism might explain the apparent incompatibility of pipolins with other integrative MGEs such as integrons.

In addition, analysis of cophylogeny between pipolins and pipolin-harboring strains showed a lack of congruence between several pipolins and their host strains, in agreement with horizontal transfer between hosts. Overall, these results indicate that pipolins can serve as a vehicle for genetic transfer among circulating *E. coli* and possibly also among other pathogenic bacteria.

## Introduction

Mobile genetic elements (MGE), comprising bacteriophages, transposons, plasmids, and insertion sequences, contribute to the great plasticity of the bacterial genome, resulting in an extremely large pangenomes that, in the case of *Escherichia coli*, can amount to more than 16,000 genes [1]. Thus, MGEs dynamics is the main source of horizontal gene transfer, which leads to the spread of antimicrobial resistance (AR) among both *E. coli* and other commensals, thereby enlarging the spectrum of resistance (the resistome) among circulating strains [2].

Pipolins constitute a recently reported new group of integrative MGEs widespread among diverse bacterial phyla and also identified in mitochondria as circular plasmids [3]. The hallmark feature of the pipolins is a gene encoding for a replicative family B DNA polymerases (PolB) with an intrinsic *de novo* primer synthesis capacity, called <u>p</u>rimer-independent PolBs (piPolBs), hence their name (<u>piPol</u>B-encoding elements). Preliminary phylogenetic analyses indicated that piPolBs would form a third major branch of PolB, besides the protein-primed PolBs (pPolBs), found in a wide range of viruses and plasmids, and RNA-primed PolBs (rPolBs), which are the principal replicative enzymes of archaea and eukaryotes, some archaea and many dsDNA viruses [3].

Because of the fact that pipolins encode the major protein required for their replication (i.e., piPolB), they are included in the proposed class of self-synthesizing (or self-replicating) MGEs, which also includes two other superfamilies of elements integrated in various cellular genomes [4]. The first superfamily comprises eukaryotic virus-like transposable elements, called Polintons (also known as Mavericks), which besides a putative pPolB, encode retrovirus-like integrases and a set of proteins hypothesized to be involved in the formation of viral particles [5-7]. The second superfamily of pPolB-encoding elements, denoted as casposons, is present in a wide range of archaea and also in a few bacteria [8]. Similar to the aforementioned self-replicating MGEs, most pipolins are integrated within bacterial chromosomes, although they are also occasionally detected as episomal plasmids. However, unlike polintons and casposons, the integrated pipolins encode for one or more integrases of the tyrosine recombinase superfamily, which could be responsible for pipolin excision and/or integration [3].

The widespread and patchy distribution of pipolins among bacteria is in agreement with an ancient origin and horizontal dispersal of this MGE group. However, pipolins from the same or related species seem closely related, as was the case for pipolins from *E. coli* [3]. The great majority of pathogenic *E. coli* strains encode for virulence-associated cassettes and antibiotic resistance genes, which are usually carried by MGEs, such as pathogenicity islands (PAIs), plasmids, integrons, etc [9-11]. However, the annotation of pipolins from *E*. coli as well as from proteobacteria did not lead to the identification of any antibiotic resistance genes or virulence factors [3].

Therefore, whereas reported evidence of mobility of polintons and casposons is limited and based on metagenomic data [12], pipolins provide the opportunity to analyze the occurrence, diversity, and dynamics of self-replicative MGEs in well-characterized commensal and pathogenic bacteria, not only in genomic or metagenomics data, but also in circulating field isolates and pathogenic variants. In this work, we surveyed the presence of pipolins in a wide collection of pathogenic strains from the Spanish *E. coli* reference laboratory (LREC). We found that pipolins, although not very abundant, are widespread among a great variety of human and animal strains belonging to different pathotypes, serotypes and sequence types (STs).

Whole-genome sequencing of the new pipolin-harboring *E. coli* strains allowed us to characterize in detail the pipolins’ hosts and also the genetic structure and phylogeny of pipolins. Most pipolins contain att-like terminal direct repeats and they are integrated in the same tRNA gene. Nevertheless, they encode a great diversity of proteins, many of which are orphans with unpredictable function. The comparison of the new strains from LREC dataset with a number of detected pathogenic *E. coli* pipolin-harboring strains from the NCBI GenBank database further confirmed our results. Finally, pangenome and cophylogeny analyses among pipolins and host strains, demonstrated that, except for strains from the same clonal complex, there is an overall lack of phylogenetic congruence between pipolins and host strains. Therefore, our results support that pipolins are a novel group of active mobile elements that might serve as a platform for horizontal gene transference among diverse pathogenic bacteria.

## RESULTS

### Limited prevalence of pipolins among *E. coli* isolates from animal and human sources

The first objective of this study was to investigate the occurrence of pipolins in *E. coli* strains causing intestinal and extraintestinal infections in humans and animals. We performed a survey of pipolin distribution among 2238 strains from the LREC collection, using a 587 nt fragment of the piPolB coding sequence as a marker. We detected 25 pipolin-harboring isolates, indicating that pipolins are not particularly abundant (1.1%) among pathogenic *E. coli* (Table 1). Interestingly, however, new pipolins are present in strains isolated from humans, swine and avian, and belonging to a wide range of pathotypes (Table 2 and Table S1).

**Table 1.** Pipolin identification survey among *E. coli* strains from diverse origins and pathotypes.

| Source | Infection | Strains | Pipolins (%) |
| --- | --- | --- | --- |
| Human | Intestinal | 608 | 8 (1.3%) |
| Human | Extraintestinal | 1061 | 9 (0.8%) |
| Swine | Intestinal | 323 | 5 (1.5%) |
| Avian | Extraintestinal | 246 | 3 (1.2%) |
| Total |  | 2238 | 25 (1.1%) |

**Table 2.**
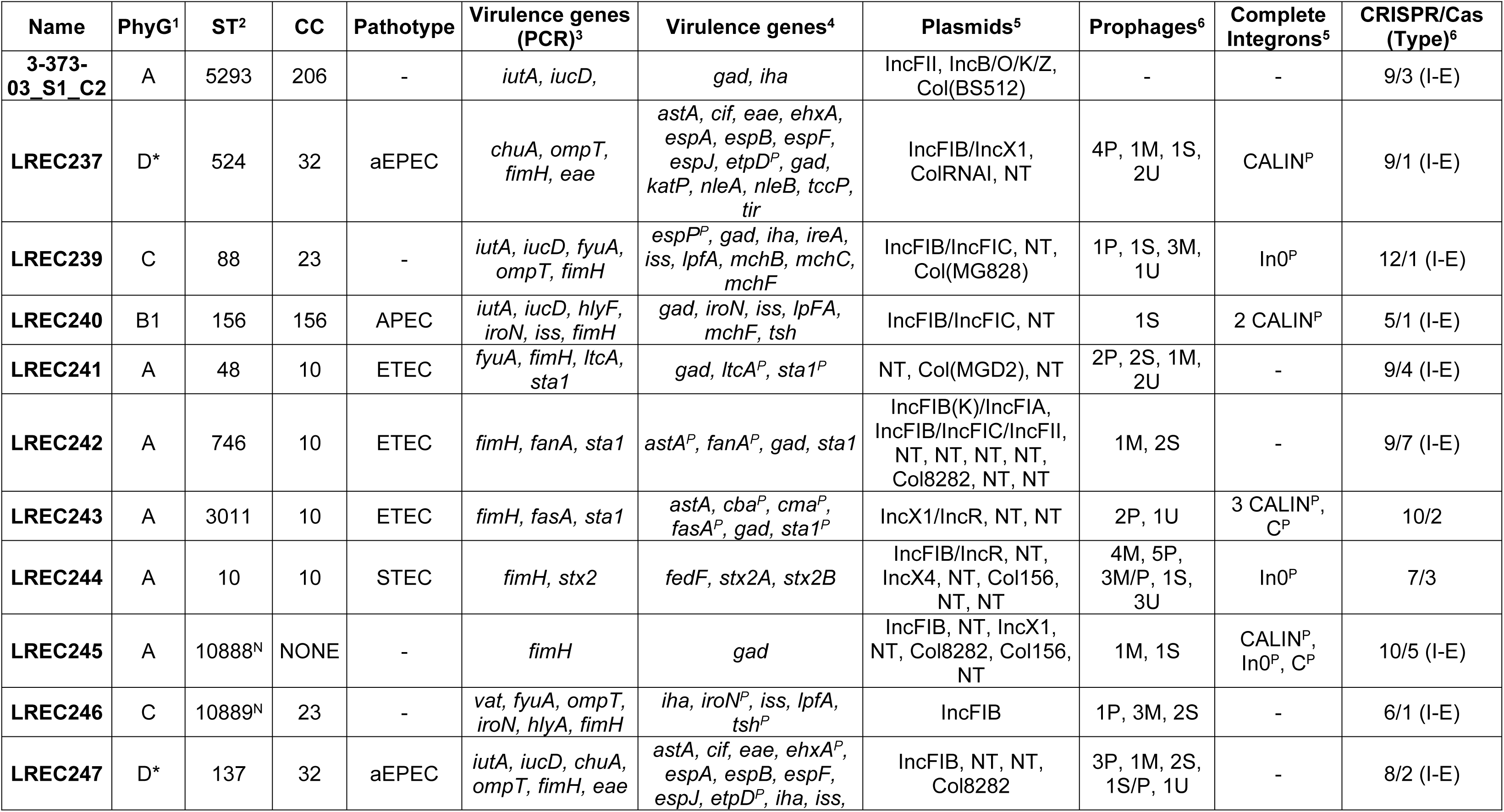
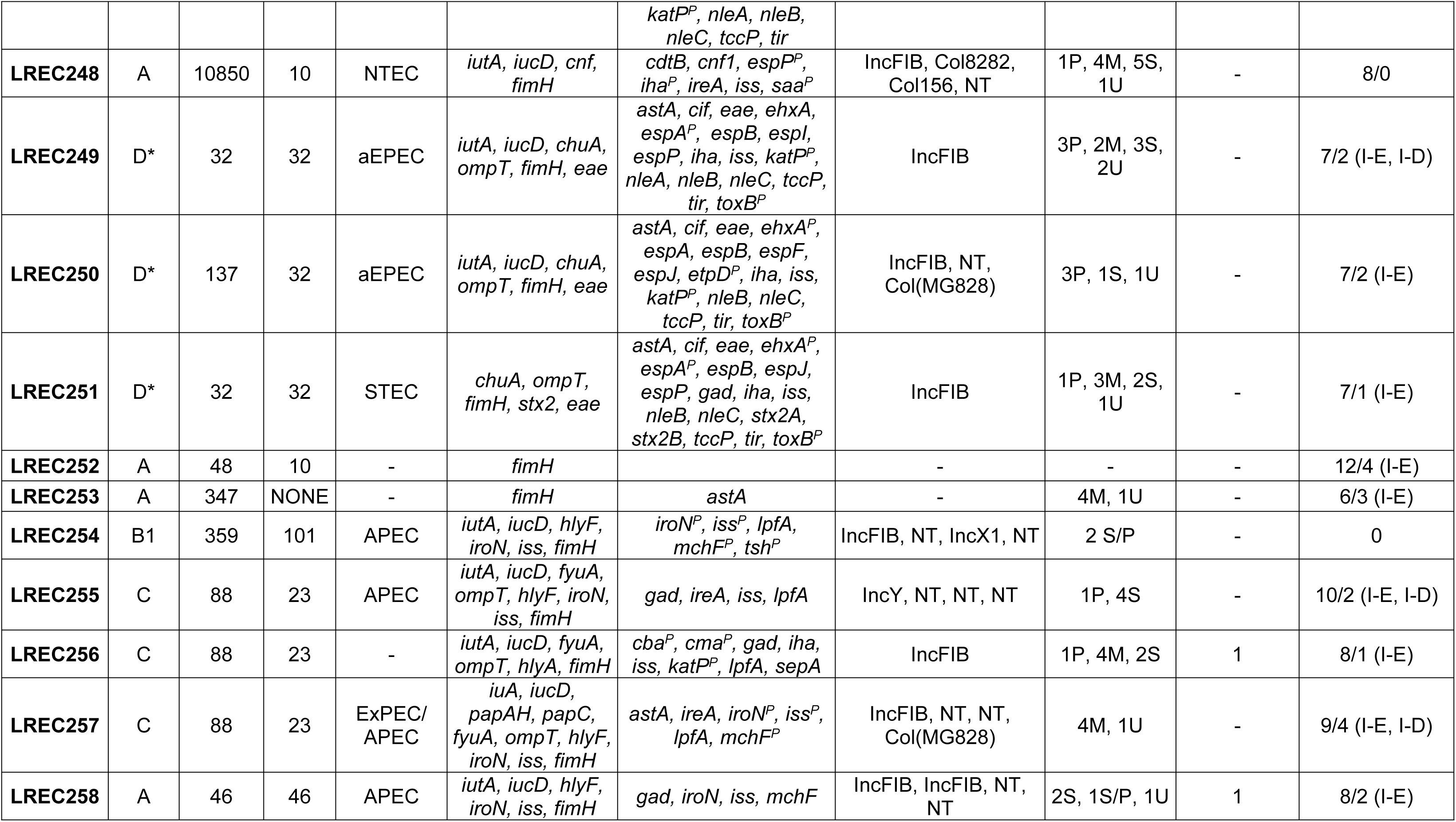
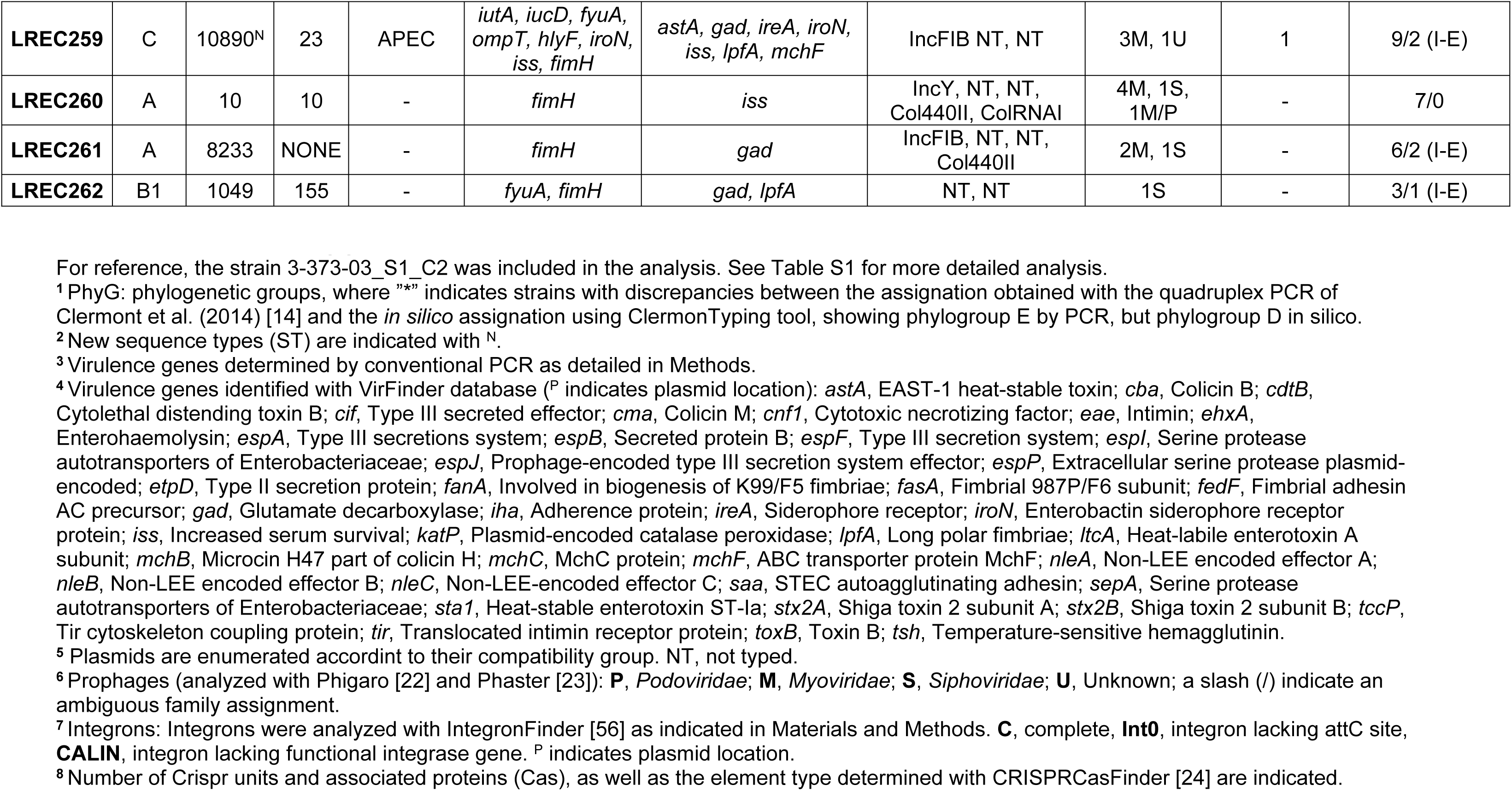
Features of the 25 pipolin-harboring *E. coli* genomes from LREC dataset.

In order to ascertain the representativity of the new pipolin-harboring strains from LREC collection, we performed a BLASTn search against the *E. coli* nucleotide database at NCBI GeneBank using the 3-373-03_S1_C2 piPolB gene as a query. This search yielded 77 hits, corresponding to piPolB-encoding ORFs or fragments from 67 *E. coli* strains, many of them scarcely characterized (Table S2). A combined phylogeny of piPolB coding sequences from LREC and GeneBank datasets (Figure S1), shows that our collection of strains carrying pipolins spans all the available diversity of *E. coli* pipolins. Therefore, the low frequency of pipolins detected in our collection is in agreement with a low prevalence of this element among circulating *E. coli* strains.

### Diversity of new pipolin-harboring *E. coli* strains

We performed a detailed characterization of LREC pipolin positive strains, both by conventional methods and whole-genome sequence (WGS) analysis (Table 2 and Table S1). We found that pipolins were present in phylogroup A (11 new strains and the reference isolate 3-373-03-S1-C2 [13]), but also in B1 (3 strains) and C (6 strains). Five strains were typed as E by quadruplex PCR typing [14] but later on reassigned as D after whole genome sequencing and *in silico* typing (see Methods for details). A similar distribution pattern was found within the GeneBank dataset, with pipolins in phylogroups A (46 strains), B1 (9 strains), C (2 strains) and D (10 strains).

The common presence of *E. coli* strains from phylogroup A in the dataset was somewhat expected, as this phylogroup is the most common among human isolates and thus very abundant in most collections [15, 16]. However, we were surprised by the absence of pipolins among B2 strains, despite the fact that this phylogroup is also very common in the LREC collection and, along with group D, it is responsible for most extraintestinal *E. coli* infections in human and animals [11, 14]. The null prevalence of pipolins among B2 strains is opposite to the pattern of occurrence of some virulence-related MGEs, such as colibactin encoding *pks* islands, which are highly prevalent in B2 and D phylogroups but were not detected in strains carrying pipolins [10, 17]. It is thus tempting to speculate whether there is an interference between *pks* islands and pipolins as they are also flanked by terminal direct repeats and found integrated into a constant tRNA [18]. Thus, the distribution of pipolins limited to phylogroups A, B1, C and D may be due to the restricted mobility beyond those groups. In line with this, these phylogenetic groups have been proposed to belong to different ancient lineages [19], downplaying a strict vertical transmission of pipolins throughout the evolutionary diversification of *E. coli* phylogroups.

Regarding multilocus sequence typing (MLST), we have detected 18 different STs among 25 LREC pipolin-harboring strains, according to the Achtman scheme [20] (Table 2), some of them are quite common in Enterobase [21], like ST88, ST10, ST46, ST48 or ST746, but also very rare varieties (ST524, ST3011, ST8233 or ST10850) and three new type sequences (ST10888, ST10889 and ST10890). Interestingly, although some of them belong to the same clonal complex (CC10), in general strains from phylogroups A and B1 span a vast diversity of STs. In contrast, LREC pipolin-harboring strains from phylogenetic groups C and D seem more homogeneous and they could be assigned to the same clonal complex (CC23 and CC32, respectively). Clear clonality among pipolin-harboring strains is more evident when we analyzed the 67 strains of GenBank dataset (Table S2), as 61.2% of the strains belong to the above-mentioned clonal complexes. Moreover, among the GenBank strains, a fourth prevalent clonal complex was CC278, observed in 5 ST278 strains of serotype O178:H7 isolated from mice.

Clonotypes and serotypes diversity is also in agreement with the presence of atypical strains (Table S1), particularly from phylogenetic groups A and B1, in which, for instance, present 14 different serotypes for 14 LREC strains. On the contrary, as in the case of STs and CCs, strains from phylogroups C and D showed overall more similar clonotypes and serotypes. Thus, the six strains of phylogroup C showed the H19 flagellar antigen and the five strains of phylogroup D showed the clonotype CH23-331 and the H28 flagellar antigen. A similar pattern can be observed among strains from the GenBank dataset (Table S2).

Analysis of virulence genes among the 25 LREC pipolin-harboring strains also allowed us to identify diverse *E. coli* pathotypes, namely, extraintestinal pathogenic (ExPEC, 1 strain), avian pathogenic (APEC, 6 strains), Shiga toxin-producing (STEC, 2 strains), enterotoxigenic (ETEC, 3 strains), atypical enterophatogenic (aEPEC, 4 strains) and necrotoxigenic (NTEC, 1 strain). However, some other common pathotypes were not detected in the pipolin-carring LREC strains, like uropathogenic *E. coli* (UPEC), typical enteropathogenic *E. coli* (tEPEC), enteroinvasive *E. coli* (EIEC) and enteroaggregative *E. coli* (EAEC).

### Antimicrobial resistance and virulence genes in LREC strains harboring pipolins

Pipolins are present in both antibiotic sensitive (10 strains) and antimicrobial resistant strains (15 strains) in the LREC collection (Table S1). Eleven strains exhibited a multidrug-resistant (MDR) phenotype. Besides, there were three extended-spectrum β-lactamase (ESBL)-producing strains, two AmpC-producing strains and two colistin-resistant strains.

In line with these results, many antimicrobial resistance genes (ARGs) were found including acquired resistance genes, point mutations and efflux/transporter genes (Table S1). We described genes conferring resistance to betalactamases (*bla*_CTX-M-15_, n=1; *bla*_CTX-M-14_, n=1; *bla*_CTX-M-1_, n=2; *bla*_TEM-1_, n=7), colistin (*mcr*-1.1, n=2), tetracycline (*tet*(A), n=6; *tet*(B), n=3; *tet*(M), n=3), aminoglycosides (*aadA1*, n=3; *aadA2*, n=4; *aadA5*, n=1; *aadA9*, n=1; *aadA13*, n=1; *ant*(3’’)-Ia, n=2; *aph*(3’)-Ia, n=2; *aph*(3’’)-Ib, n=5; *aph*(4)-Ia, n=2; *aph*(6)-Id, n=4; *aac*(3)-IV, n=2 and *aac*(3)-IIa, n=1), phenicols (*catA1*, n=5 and *cmlA1*, n=3), trimethoprim (*dfrA1*,n=2 and *dfrA12*, n=3), lincosamides (*lnu*(F), n=2), macrolides (*mph*(A),n=1; *mph*(B),n=1 and *mef*(B), n=1), quinolones (*qnr*S1, n=1 and *qnr*B19, n=1) and sulfonamides (*sul*1, n=3; *sul*2, n=2 and *sul*3, n=3). Furthermore, we found chromosomally encoded point mutations in the *gyrA* and *parC* genes conferring resistance to quinolones in nine strains and in the *ampC* promoter conferring resistance to betalactamases in three strains.

In summary, LREC pipolins -carrying strains, harbor a repertoire of ARGs, as expected of pathogenic *E. coli* strains, ruling out any correlation among ARGs and pipolins, in line with the diversity of phylogenetic groups, STs and pathotypes they belong to.

### Pangenome and mobilome of *E. coli* strains harboring pipolins

We assembled the pangenome of the 25 LREC strains carrying pipolins plus the reference strain, 3-373-03_S1_C2 with Roary [26], resulting in 10,178 different genes (Figure S2). Among those, 2,998 genes (29.45%) corresponded to the core-genome and were present in all strains, 327 were soft-core genes present in 95-99% of genomes and 2,259 were shell genes present in 15-95% genomes. As expected from such a diverse pangenome, almost half of the genes (4594, 45.13%) were cloud-genes, found in less than 15% of strains. This diversity is even more evident when the pangenome of both (LREC and GenBank) datasets are analyzed together, with a total of 16,675 genes, only 934 genes comprise a core-genome and more than two-thirds of the genes in the cloud-genome (11,175, 67%). As such, the number of both total and unique genes associated with the cloud gene set increased consistently with the number of genomes (Figure S2, B-D). In conclusion, notwithstanding the clonality of several strains, pangenome analysis indicates that pipolins are present in a wide variety of *E. coli* strains.

Pangenome analysis and core-genome based phylogeny reconstruction of the LREC strains (Figure 1), clustered pipolin-harboring strains in agreement with the assigned phylogenetic groups and clonal complexes, and this congruence is maintained for the phylogeny inferred from the core genome of all analyzed strains carrying pipolins (Figure S3). Similar results were obtained when the phylogeny of the strains was constructed by single-nucleotide polymorphism in EnteroBase [21], although some of the strains are not available in this database (Figure S4).

**Figure 1.**
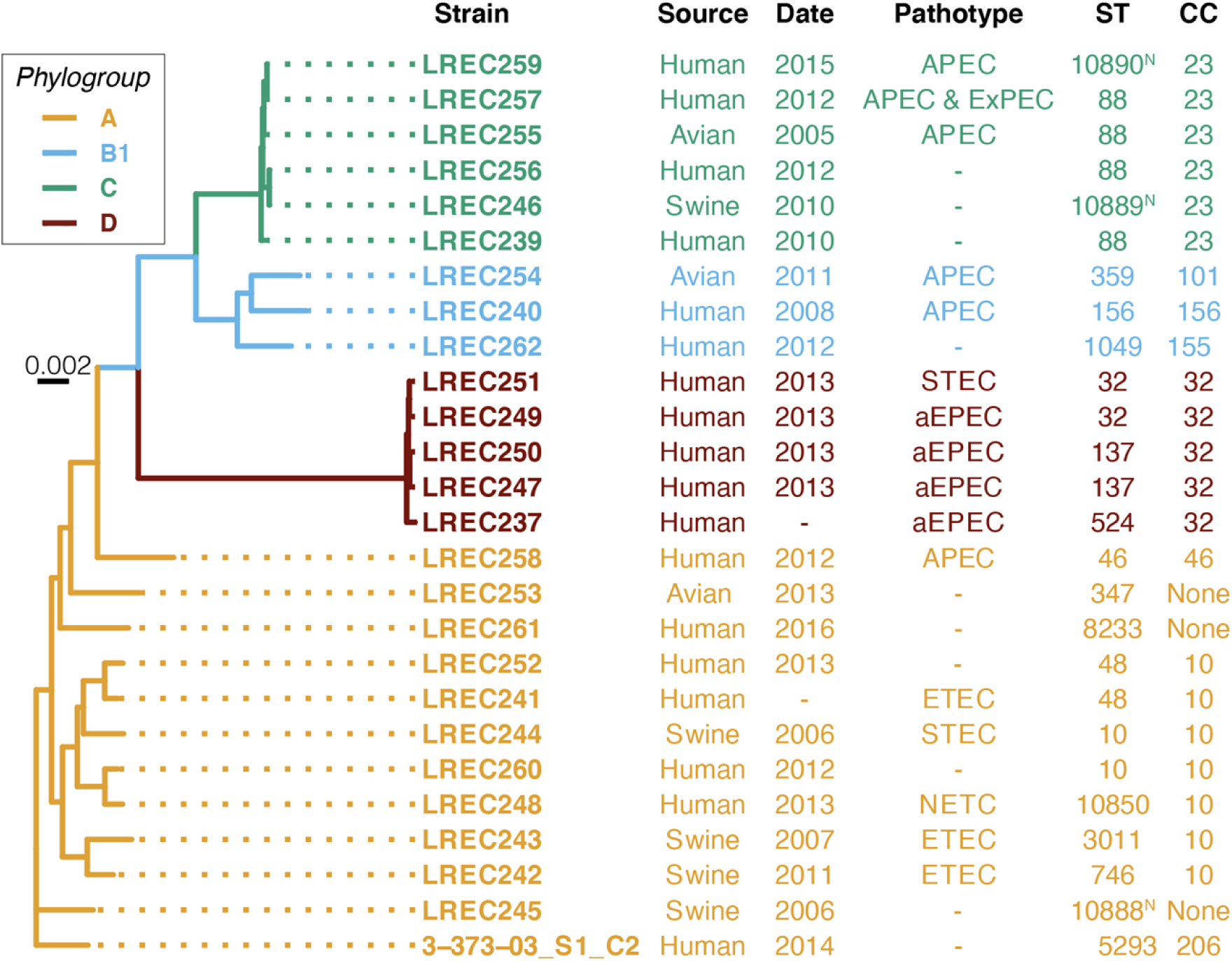
Maximum-likelihood tree generated from the core-genome data of new pipolin-harboring strains. Strain names are colored according to the phylogenetic groups as indicated. Previously described pipolin-harboring isolate 3-373-03_S1_C2 was included as a reference. The best-fit model was GTR+F+R2 for all considered criteria in ModelFinder [25]. Scale bar indicates substitution rate per site. The main features are indicated on the right: source, isolation date, pathotype, sequence type (ST) and clonal complex (CC). New ST combinations assigned at Enterobase are indicated with ^N^.

Disregarding pipolins, when the mobilome of our strains was analyzed, we could identify the typical variety of plasmids and other mobile elements. PLACNETw [27] allowed us to identify and assemble several plasmids in most of the strains (Table 2 and Table S1) and 79 elements could be extracted, which would correspond to at least a total of 86 plasmids, as some of the elements contained markers from more than one incompatibility group. All strains carried a least one plasmid moving in a range from 1 to 9, except for strains LREC252 and LREC253 (Table 2 and Table S1). Plasmids from IncF incompatibility group were the more prevalent including IncFIB (n=20), IncFIA (n=1), IncFII (n=2) and IncFIC (n=3) replicons. Followed by Col-like plasmids including Col8282 (n=4), Col(MGD2) (n=1), Col(BS512) (n=1), Col(MG828) (n=3), Col156(n=3), Col440II(n=2), and ColRNAI (n=2). However, we also described the presence of other incompatibility groups like IncX1 (n=3), IncY (n=2), IncR (n=2), IncX4 (n=1) and IncB/O/K/Z (n=1). Furthermore, 42 cryptic plasmids could not be affiliated with any category as they lack any known replication origin. Nonetheless, none of those correspond with an episomic pipolin and indeed piPolB-containing contig was detected as a single copy portion of the chromosome, which suggests that pipolin excision is negligible under standard growth conditions or undetectable by Illumina sequencing. As expected, many of the reported antimicrobial and virulence factors were plasmid-borne genes, since conjugative plasmids, along with other MGEs, are the most successful genetic platforms allowing the horizontal transfer of antimicrobial resistance and virulence determinants among pathogenic *E. coli* isolates [11, 28, 29].

Most of the strains contained genes from one or more prophages and, as expected, sequence arrays and genes from the CRISPR/Cas immunity system (Table 2).

Strikingly, integrons seem quite uncommon, as we could detect complete integrons in 5 out of 25 new strains (20%) and only 5 hits were detected in the pipolin-harboring genomes from Genbank (8.9%), whereas they are often reported to be usually highly prevalent and present in more than half of pathogenic *E. coli* strains [30-33].

### Mapping and extraction of new pipolins from LREC and GenBank datasets

Besides the presence of a piPolB gene, pipolins are characterized for the presence of att-like terminal direct repeats that might be involved in recombination-mediated excision/insertion, often in a tRNA site [3]. We extracted pipolins from both, LREC and GenBank pipolin-harboring strains, using a custom bioinformatics pipeline that entailed searching for piPolB gene or its gene fragments and terminal direct repeats to determine the element bounds (see Methods for details). Except for the strain LREC243, two att-repeats could be detected in all genomes. In the cases when att-repeats were located on the same contig or on a complete chromosome, the piPolB always sitting within the repeats, which confirmed the basic structure of all *E. coli* pipolins (Figure 2 and Figure S5). This structure could be reconstructed also when piPolB and att-repeats were not on the same contig (see Methods). Reparably, all *E. coli* pipolins are integrated in the same point, at the Leu-tRNA gene, except for the pipolin from LREC252 strain that looks inconsistent with other pipolins. The att repeat that overlaps with the tRNA gene was represented and denoted as right end (attR). In some genomes, three att-repeats were detected, as those pipolins seem to share the integration site and mechanism with some prophage, as previously detected for the enterotoxigenic *Escherichia coli* H10407 strain [3]. Indeed, comparison of the genetic structure of all pipolins (Figure S5), confirmed that a similar myovirus enterophage is present next to pipolins from eight strains, spanning phylogroups A (H10407, 2014EL-1346-6 and 99-3165), C (LREC239 and LREC246) and D (112648, 122715, 2015C-3125 and FWSEC0002). In addition, the presence of transposases and associated genes indicates that genetic islands and insertion sequences can as well contribute to the variability of pipolins, particularly in the case of stains LREC248 and LREC252, expanding also the pipolin gene repertoire (see below). The cohabitation of casposons with other MGEs is also common [12], although in that case, the associated element seems to provide a vehicle for horizontal transfer. On the contrary, as pipolins possess att-like direct repeats and one or more recombinases (see below), they can be considered as self-transferable.

**Figure 2.**
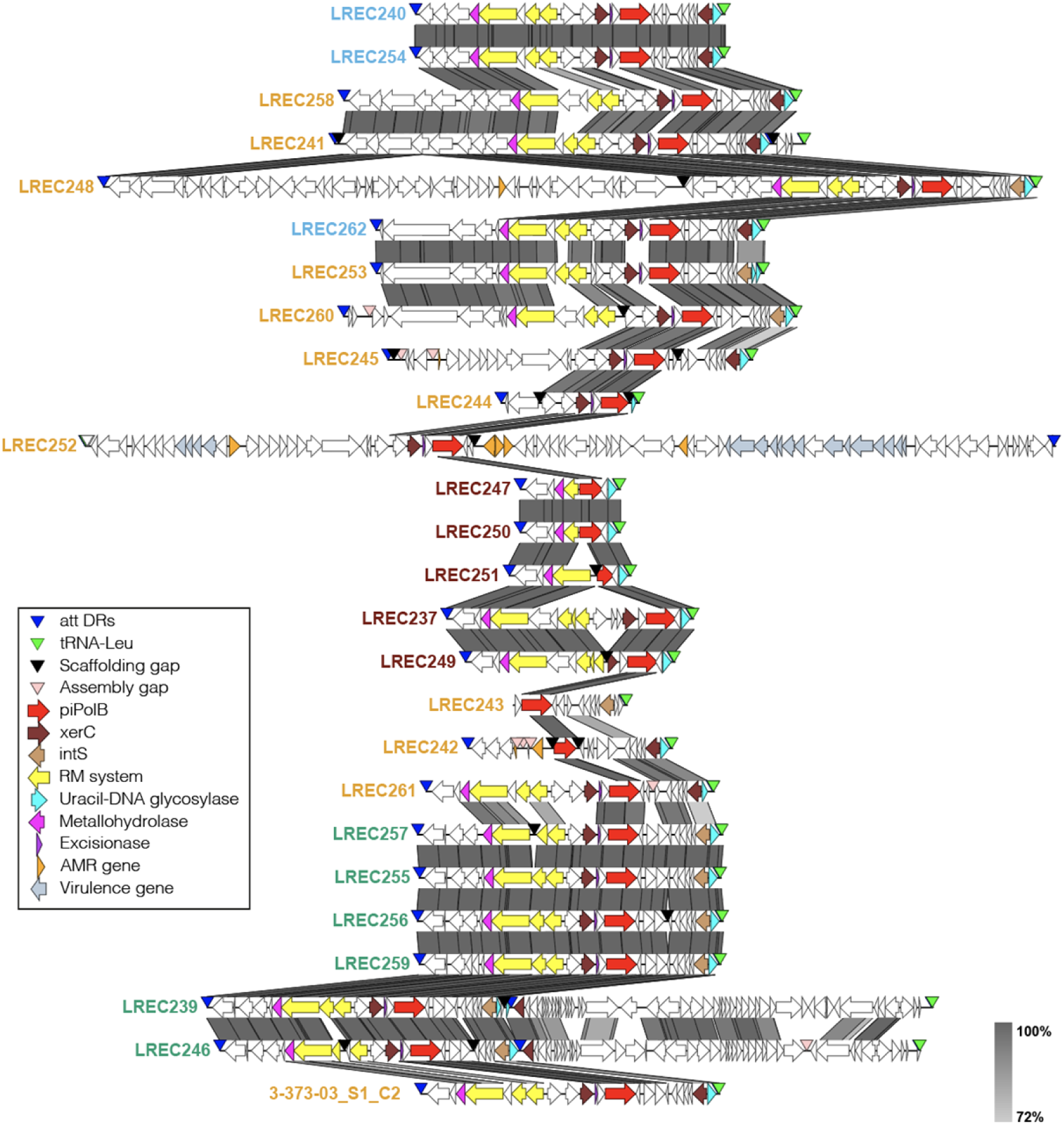
Genetic structure of new pipolins from LREC collection. Predicted protein-coding genes are represented by arrows, indicating the direction of transcription and colored following Prokka annotation as indicated in the legend. The greyscale on the right reflects the percent of amino acid identity between pairs of sequences. The image was generated by EasyFig software and re-annotated pipolins sorted according to the hierarchical clustering of the gene presence/absence matrix. Names of pipolin carrying strains are colored according to phylogroups as in Figure 1.

Altogether, mapping and extraction of the new *E. coli* pipolins, confirmed that, despite a great diversity, they share basic genetic structure and they can likely be mobilized using the same mechanism.

### Pipolins annotation and pangenome analysis

The 92 extracted pipolins were reannotated with a custom pipeline using Prokka [34] (see Methods), followed by Roary [26] for pangenome analysis of *E. coli* pipolins, identifying a total of 272 genes. Remarkably, the core- and soft-core genomes are made up of a single gene cluster, the piPolB, and a XerC-like tyrosine-recombinase, respectively. In line with this, the shell genome contains only 38 genes, whereas 232 genes (85%) are cloud-genes, present in less than 15% of pipolins. Despite the great variety of different genes, some groups of pipolins share a similar gene composition and, as about 75% of genes are provided by about one-third of the pipolins (Figure S5). Although a certain level of synteny and modular organization can be detected (Figure 2, Figure S5), genetic rearrangements, including inversions, duplications, and deletions, which often lead to gene exchange, are also frequent, as well as truncations and disruptions. Even truncated forms of piPolBs or XerC-like recombinases can also be detected, which might lead to impairment of replication or mobilization of pipolins. Overall, the genetic repertoire and structure of analyzed pipolins suggests that they can exchange genetic information among *E. coli* strains.

A detailed functional analysis of shell core genes is shown in Table S3. As mentioned above, besides piPolB, pipolins very often include one or more XerC and IntS (bacteriophage-type) tyrosine recombinases. When two complete recombinase genes are present, one of them is always close to an excisionase-like protein. A type-4 Uracil DNA glycosylase is also very frequent. Other proteins with DNA binding domains like mobilization proteins as well as components of restriction-modification systems are also common. Very few antimicrobial resistance genes or virulence genes are detected, always associated with associated MGEs (prophages or transposons).

In summary, a pipolin basic unit is composed of direct terminal repeats encompassing a piPolB gene and a variety of genes, most of them related to the metabolism of nucleic acids.

### Cophylogeny of pipolins and host strains suggest pipolins horizontal transfer

Since the presence of the piPolB gene is the hallmark of pipolins and it constitute the only core gene, we performed a phylogenetic analysis of the new piPolB sequences from the new pipolin-harboring *E. coli* strains. Although some of the new annotated piPolB genes are partially truncated, particularly those from pipolins in phylogroup D strains, they have a high degree of identity, above 98.8% in the aligned regions. Phylogeny of the LREC piPolBs (Figure 3) underlined again the similarity among pipolins in clonal strains that belong to phylogroups C and D, but sequences from phylogroups B1 and A were mixed together. A somewhat similar pattern was obtained for the phylogenies of XerC-like recombinases (Xer_C_2 group from Roary, see Table S3) and UDGs in the combined collection of pipolins (Figure S6). In order to assess the significance of different phylogenetic trees, we calculated the cophenetic correlation coefficient (CCC,[35]) among them as indicative of phylogenies clustering congruence. When comparing the piPolB and XerC phylogenies, the CCC was 0.29; for the comparison of piPolB and UDGs phylogenies it was 0.38; and a value of 0.27 was obtained for UDGs vs XerC comparison, indicating very low clustering similarity among different pipolins genes. Thus, increasing the number of genes in pipolins phylogeny, would increase the noise, to the detriment accuracy. Therefore, we considered only the phylogeny of piPolB coding sequence, as the only hallmark gene for all pipolins, for subsequent cophylogeny analyses. This can be considered also as a functional criterion, since piPolB is probably essential for episomal pipolin replication, being thus in agreement with conventional taxonomy of plasmids or other elements from the prokaryotic mobilome [36, 37].

**Figure 3.**
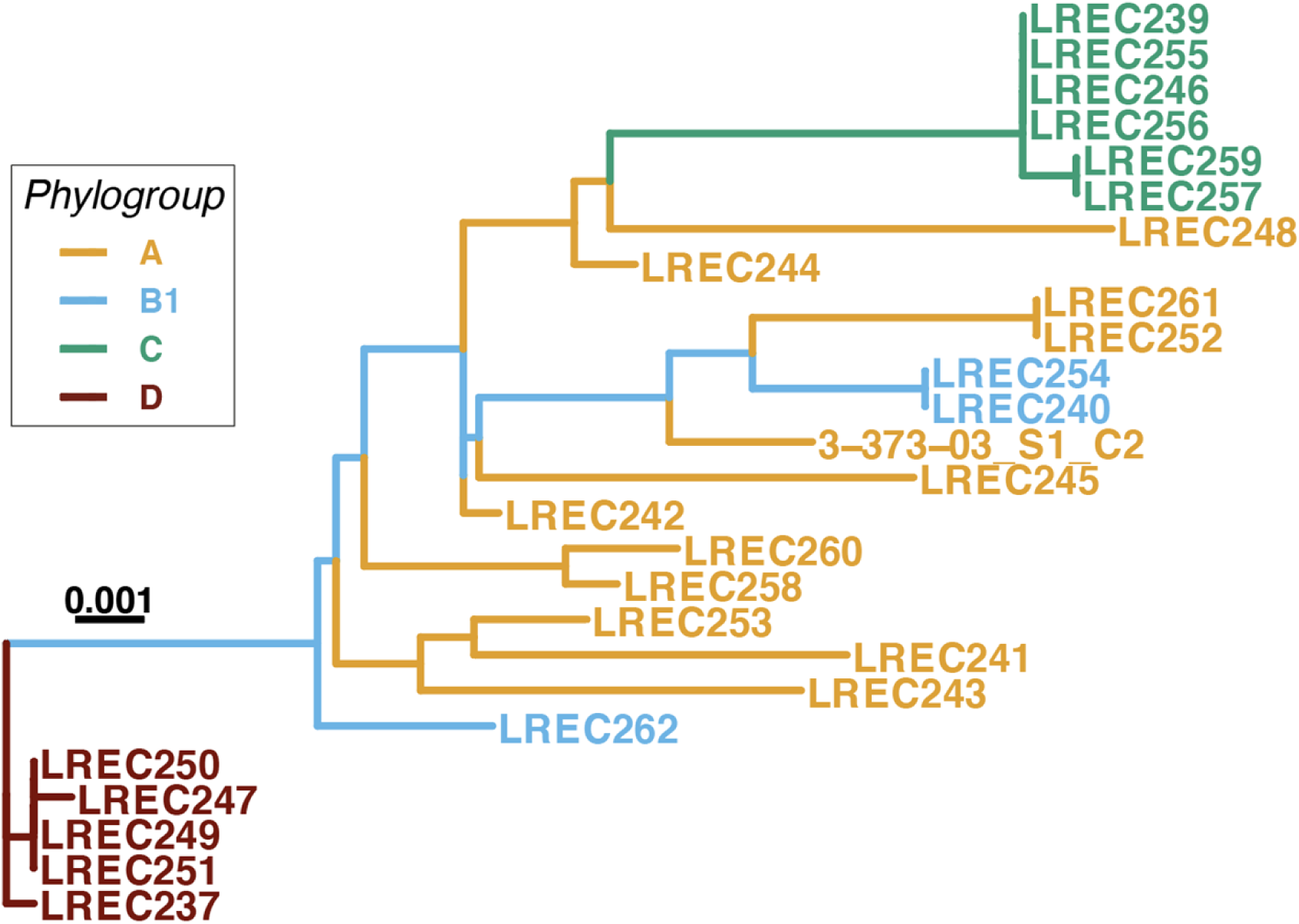
Maximum-likelihood tree of the new piPolB genes from the LREC dataset. As indicated, strain names are colored based on the phylogenetic group of strains. Previously described pipolin-harboring isolate 3-373-03_S1_C2 was included as a reference. The best-fit model was GTR+F+R2 for all considered criteria by ModelFinder [25]. Scale bar indicates the substitution rate per site.

The tanglegram in Figure 4 allows us to visualize the cophylogeny between piPolBs and *E. coli* strains carrying pipolins. This plot reveals a complex association pattern, with numerous crisscrossing lines that suggest incongruence between the two phylogenies and, in line with this, the CCC is quite low, 0.14.

**Figure 4.**
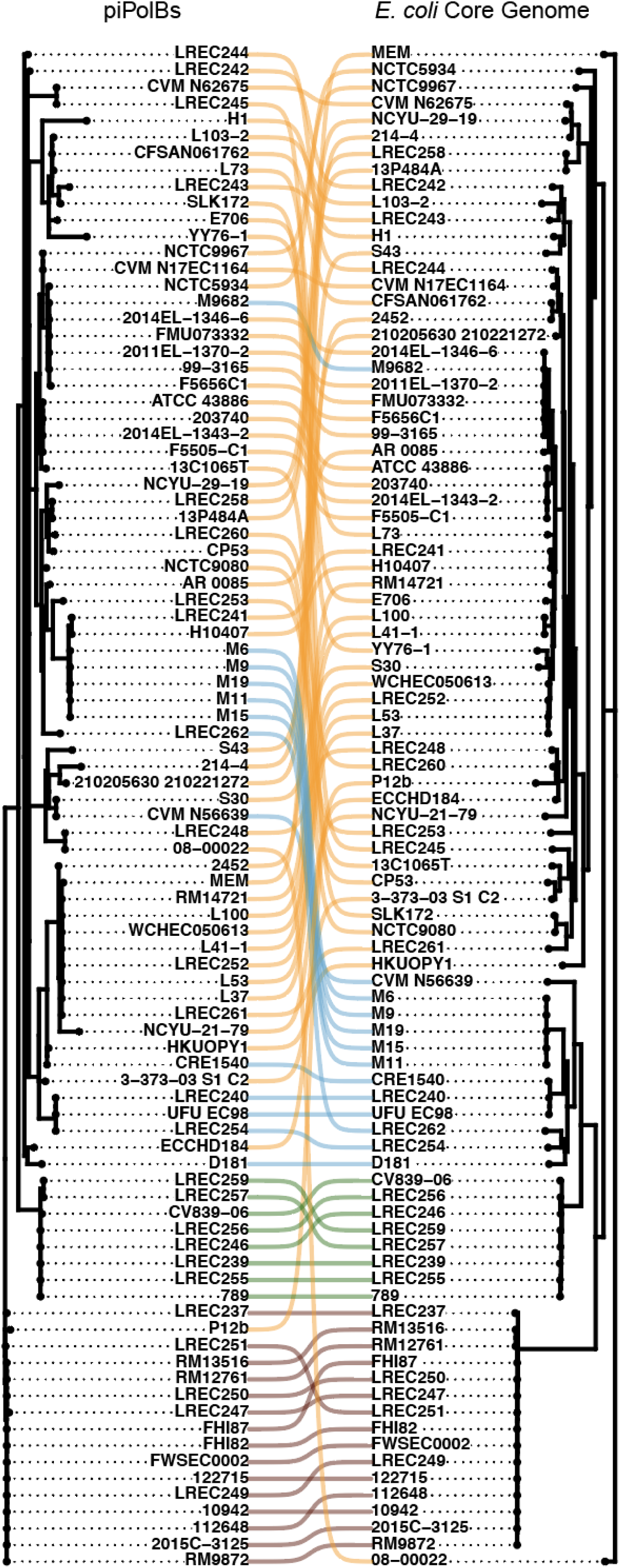
Cophylogeny of pipolins and host strains. Tanglegram representation of maximum-likelihood comparative phylogenies of piPolB and host strains core genome as hallmark of pipolins. Modelfinder Best-fit models were K3Pu+F+R2 and GTR+F+R7, respectively. Compared phylogenies are also displayed in Figures S1 and S3, respectively. Links between pipolins and *E. coli* strains are clored based on the phylogenetic groups in Figure 3.

Furthermore, we tested the cophylogeny between this group of pipolins and the host strains using PACo (Procrustean Approach to Cophylogeny) [38]. This method considers both clustering and relative distances, by using a procrustean approach in distance-based statistical shape analysis of phylogenetic trees that provide global-fit values (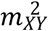) for the trees’ shape comparison. These analyses allow the detailed characterization of parasite-host and virus-host evolutionary interactions, under the null hypothesis that the topology of the host tree cannot predict the topology of the parasite tree [39-41]. The 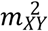 are inversely proportional to the topological congruence between the two phylogenetic trees [38]. In our case, the 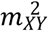 is 0.017, with a p-value of 0.46. Thus, in agreement with the very low CCC value, we cannot reject the null hypothesis, indicating that a significant portion of the pipolins tree topology does not depend on (i.e., cannot be predicted by) the host strains phylogeny. Moreover, when we analyzed the cophylogeny of pipolins and strains from each phylogenetic groups, we found that the Procrustes residuals of the pipolins and strains from phylogroup C and D, but not those from groups A or B1 (Figure S7A-D), were significantly smaller than the remainder of the interactions in the cophylogeny network, indicating that these interactions show significantly greater phylogenetic congruence than the rest. These lineages reflect the tree topologies of their host strains, indicating either co-evolutionary association or restricted horizontal transfer to highly related strains. Interestingly, some of the pipolins from strains from phylogroup D seem to have a truncated form of XerC_2 gene that only contains the Arm DNA binding domain present (see Figure 3 and Figure S5, LREC247, LREC250 and LREC251), which could explain the cophylogeny of those pipolins with their host strains (i.e. low horizontal transfer), as they seem to lack any recombinase/integrase activity.

Further, to identify the contribution of each *E. coli* strain to the overall cophylogeny structure, we evaluated the Jack-knifed squared residual values (Figure S7E). This analysis showed that an important proportion of pipolins from phylogroups C and D, but also for a few of the strains in phylogroup A, like L103-02, F5005-C1, ATCC-43886 or LREC258, among others, present low squared residual values, indicating congruent topologies between piPolBs and genomic trees. Interestingly, all the pipolins that present a tandem cohabitation with a prophage are within this group, in line with the previously hypothesized inactivation of pipolins as a consequence of the prophage insertion. On the other hand, pipolins with high squared residual values from phylogroups A and C would not have been evolutionary linked to their host strains, in agreement with the previous results.

Overall, we can conclude that the pipolins diversity is poorly congruent with the strains phylogeny and their distribution is rather indicative of a patchy distribution amongst a wide variety of pathogenic *E. coli* strains, as expected from horizontally transferred MGEs. This pattern may reflect the wide distribution of pipolins beyond *E. coli*, dispersed among major bacterial phyla, namely Actinobacteria, Firmicutes, and Proteobacteria, as well as in mitochondria [3].

### Conclusion and Perspectives

Self-replicating integrative MGEs are highly diverse members of the prokaryotic and eukaryotic mobilome that seem to be involved in key evolutionary events, including the origin of the CRISPR-Cas systems from casposons and the evolution of several groups of eukaryotic dsDNA viruses from polintons. However, the information about their mobilization is limited and biased by the availability of genomic and metagenomic data in databases.

Pipolins represent a unique group of recently discovered, self-replicating integrative elements, which display broad distribution among bacteria. Their presence in mitochondria and phylogenetic analysis suggested ancient origin [3], though their evolutionary history remains unclear. Here we have undertaken the first characterization of a predicted self-replicating MGE within a collection of circulating human and animal pathogenic strains of *E. coli*.

Despite the relatively low frequency of pipolins, our results confirmed that, rather than correlating with a certain phylogenetic group or pathotype, or the presence of a particular antimicrobial resistance, pipolins show a patchy distribution among a variety of circulating *E. coli* strains, both from a collection screening and a GenBank survey. We could detect clonality of several groups of isolates carrying pipolins, belonging to the C and D phylogenetic groups. Moreover, their clonality reflects the phylogeny of their harbored pipolins, indicating that their mobility cannot be detected. However, mobilization of some of those pipolins seems impaired by the inactivation of XerC recombinase. On the contrary, the phylogeny of more diverse and numerous hosts from phylogroups A and B1 shows a lack of congruence with their pipolins, ruling out a monophyletic origin or strict vertical transmission of pipolins.

Analysis of the genetic structure and pangenome of pipolins from *E. coli* showed that they are more dynamic and flexible mobile elements than might be foreseeable from previous work. Pipolins encode a great diversity of genes, with an average of more than 7 different genes per element, with piPolB being the only shared gene in all pipolins. However, virulence and antimicrobial resistance genes were not detected in pipolins in our dataset, which may explain their low prevalence in pathogenic *E. coli* strains, but also raise questions about their biological role and evolution. Moreover, whereas diversity of plasmids is usual for pathogenic strains, our results suggest a possible interference between pipolins and other integrative MGEs, like *pks* islands or integrons, usually highly prevalent among pathogenic strains of Gram-negative bacteria.

Altogether, our results provide evidence for horizontal transfer of pipolins and suggest that pipolins are an active platform for horizontal gene transfer in *E. coli*, and they also pave the way for further analysis in other clinically relevant bacteria with pipolins.

## Methods

### Pipolin screening among *E. coli* strains

A total of 2238 *E. coli* strains causing intestinal and extraintestinal infections in humans and animals from LREC collection were tested for the presence of pipolins by PCR using primers piPolB_FW (5’-GTTTTTTGACAAATTGCCCACTTG) and piPolB_RV (5’-CATATCAGAAAACACCGTCCG).

### Conventional typing of LREC pipolin-harboring strains

The 25 LREC pipolin-harboring strains were first characterized by conventional typing. The determination of O and H antigens was carried out using the method previously described by Guinée et al. [42] with all available O (O1 to O181) and H (H1 to H56) antisera. Isolates that did not react with any antisera were classified as O non-typeable (ONT) or HNT and those non motile were denoted as HNM. Assignment to the main phylogroups (A, B1, B2, C, D, E, F) was based on the PCR protocol of Clermont et al. [43]. The sequence types (STs) were established following the multilocus sequence typing (MLST) scheme of Achtman by gene amplification and sequencing of the seven housekeeping genes (*adk, fumC, gyrB, icd, mdh, purA*, and *recA*) according to the protocol and primers specified at the E. coli MLST web site (http://mlst.warwick.ac.uk/mlst/dbs/Ecoli) [20]. Clonotype identification was determined by *fumC* and *fimH* (CH) sequencing [44].

Virulence factor (VF)-encoding genes of *E. coli* causing intestinal and extraintestinal infections were screened by PCR [45, 46]. The virulence gene score was the number of virulence-associated genes detected. The isolates were designed presumptively as extraintestinal pathogenic *E. coli* (ExPEC) if positive for ≥2 of 5 markers, including *papAH* and/or *papC, sfa/focDE, afa/draBC, kpsM II*, and *iutA* [47], as uropathogenic *E. coli* (UPEC) if positive for ≥3 of 4 markers, including *chuA, fyuA, vat*, and *yfcV* [48], as avian pathogenic *E. coli* (APEC) [49] if positive for ≥ 4 of 5 markers (*hlyF, iutA, iroN, iss* and *ompT*), and as necrotoxigenic *E. coli* (NTEC) if positive for *cnf1, cnf2* or *cnf3* genes [50]. In addition ten VF-encoding genes specific for pathotypes of diarrheagenic *E. coli* (DEC) were screened by PCR and the strains were designed as: typical enteropathogenic *E. coli* (tEPEC) (*eae*+, *bfpA*+, *stx*_1_-,*stx*_2_-), atypical enteropathogenic *E. coli* (aEPEC) (eae+, *bfpA*-, *stx*_1_-,*stx*_2_-), Shiga toxin-producing *E. coli* (STEC) (*stx*_1_+ and/or *stx*_2_+), enterotoxigenic *E. coli* (ETEC) (*eltA+* and/or *est*+), enteroinvasive *E. coli* (EIEC) (*ipaH+*), and enteroaggregative *E. coli* (EAEC) (*aatA+, aaiC+* and/or *aggR+* [46].

### Antimicrobial susceptibility screening of LREC pipolin-harboring strains

Antimicrobial susceptibility was determined by minimal inhibitory concentrations (MICs). Resistance was interpreted based on the recommended breakpoints of the CLSI [51]. Thirteen classes of antimicrobial agents were analyzed: penicillins (ampicillin, AMP), penicillins and β-lactamase inhibitors (amoxicillin-clavulanic acid, AMC; piperacillin-tazobactam, PTZ), non-extended spectrum 1st and 2nd generation cephalosporins (cefalotin, KF; cefazolin, CFZ; cefuroxime, CXM), extended-spectrum 3 rd and 4 th generation cephalosporins (cefotaxime, CTX; ceftazidime, CAZ; cefepime, FEP), cephalosporins and β-lactamase inhibitors (cefotaxime and clavulanic acid, CTXc; ceftazidime and clavulanic acid, CAZc), cephamycins (cefoxitin, FOX), carbapenems (imipenem, IMP; ertapenem, ETP), aminoglycosides (gentamicin, GEN; tobramycin, TOB), nitrofurans (nitrofurantoin, F), quinolones (nalidixic acid, NAL; norfloxacin, NOR; ciprofloxacin, CIP), folate pathway inhibitors (trimethoprim-sulphamethoxazole, SXT), phosphonic acids (Fosfomycin, FOS) and polymyxins (colistin, CL). *E. coli* multidrug resistant (MDR) was defined as resistance to one or more agents in three or more classes of tested drugs [52].

### Whole Genome Sequence (WGS) and *in silico* characterization of *E. coli* strains carrying pipolins

WGS was carried out in an Illumina HiSeq1500 (2×100 or 2×150 bp) following standard protocols. Briefly, libraries for sequencing were prepared following the TruSeq Illumina PCR-Free protocol. Mechanical DNA fragmentation was performed with Covaris E220, and the final quality of the libraries assessed with Fragment Analyzer (Std. Sens. NGS Fragment Analysis kit 1-6000 bp). The libraries were then sequenced, and reads were trimmed (Trim Galore 0.5.0) and filtered according to quality criteria (FastQC 0.11.7). Obtained sequences are available as a NCBI Bioproject PRJNA610160 (see Table S1 for Biosamples Ids and Enterobase Uberstrain codes for each strain). Strain 3-373-03_S2_C2 [13] was sequenced and analyzed in parallel as a reference, but it was not included in the BioProject to avoid redundancy.

The reconstruction of the genomes and plasmids in the genomes was carried out using the methodology PLAsmid Constellation NETwork (PLACNETw) [27]. The assembled contigs, with genomic size ranging between 4.5 and 5.51 Mbp (mean size 5.08 Mbp), were annotated by Prokka [34]. Predicted CDS were analyzed using ABRicate [53] for the presence of antibiotic resistance (ResFinder V2.1.), virulence genes (VirulenceFinder v1.5), plasmid replicon types (PlasmidFinder 1.3./PMLST 1.4.), and identification of clonotypes (CHTyper 1.0), sequence types (MLST 2.0) and serotypes (SerotypeFinder 2.0). PointFinder V3.2 was used in order to find antibiotic resistances encoded by chromosomal mutations (90% min. ID and 60% min. length thresholds) [54]. Phylogroups were predicted using the ClermonTyping online tool [55]. Moreover, to characterize the strains mobilome, prophages were searched with Phigaro [22] and Phaster [23]; CRISPRCasFinder [24] was used for the report of CRISPR/Cas cassettes and a custom database of *IntI1, Intl2, Intl3, qacEdelta1* and *sul1* genes was used for identification of integrons and subsequent integrity analysis with IntegronFinder [56]. For Pipolins-harboring strains from GenBank (see below), the presence of integrons was also analyzed by the same method using the chromosome sequence and then in Integrall database ([57], updated on 1 April 2020), which allowed us the identification of one more integron, located in a plasmid. All predictions were called applying a select threshold for identification and a minimum length of 95 and 80%, respectively. Pangenome analysis was performed with Roary [26], which generated a codon aware alignment using Prank [58]. This alignment was then used for best-fit maximum likelihood-phylogenetic construction of phylogenetic tree IQTree Modelfinder [25]. For reference, single Nucleotide Polymorphism(SNP) tree were performed with EnteroBase [21], which runs a number of pipeline jobs with The Calculation Engine (TCE) in the order refMasker, refMapper, refMapper_matrix and matrix_phylogeny.

### Pipolins extraction and re-annotation

Pipolin-harboring *E. coli* genomes were retrieved from NCBI Genbank nucleotide collection using Blast [59] (October 30, 2019) and the piPolB coding sequence *from E. coli* 3-373-03_S1_C2 (GenBank id. NZ_JNMI01000006, 80192-82792) as a query. Highly similar piPolBs from related *Enterobacteriaceae* (*Citrobacter* sp., *Enterobacter* sp., and *Metakosakonia* sp.) were also detected but we discarded them to facilitate the analysis. In total, 92 *E. coli* genomes (25 from LREC collection and 67 from GenBank) were employed in the subsequent analysis.

Pipolins from the obtained genomes were extracted for detailed characterization using a custom pipeline detailed as follows. Pipolin boundaries can be defined by att-like terminal direct repeats [3]. Based on that, the nucleotide BLAST was performed using one of the att-repeat sequences from the 3-373-03_S1_C2 isolate as a query against each of 92 *E. coli* genomes. In some cases, att-repeats and piPolB were located on different contigs, posing a challenge for us to understand the order and orientation of the contigs which parts of the contigs belong to a pipolin. We assumed that att-repeats should be headed in the same direction as they are direct repeats and that one of them could overlap with a tRNA gene on the opposite strand. For consistency, we referred to the latter att-repeat as attR and expected it always to be the rightmost att. According to these assumptions, we scaffolded the disrupted pipolin regions into a continuous sequence using a custom Python script. During scaffolding, different parts of a pipolin region were connected up by introducing the “assembly_gap” feature key of unknown length (DDBJ/ENA/GenBank Feature Table Definition, Version 10.9 November 2019). Comparative representation of the genetic structure of pipolins was generated by Easyfig [60].

The extracted and scaffolded pipolin sequences were re-annotated by the Prokka pipeline [34]. This pipeline allows usage of different databases for protein annotation, among those we have been using Bacteria-specific UniProt (updated 16.10.2019), HAMAP (updated 16.10.2019) and Pfam-A (updated 08.2018). After the first try, ∼50 % of pipolin ORFs left unannotated and were classified as “hypothetical proteins”. Since the pipolins annotation was quite incomplete, we attempted to improve the annotation of pipolin genes using HHpred [61] for the most common pipolin ORFs, as defined by the Roary analysis. We considered the found hits as homologous if 1) the probability was >90 %, 2) E-value <0.01, 3) secondary structure similarity was along the whole protein length, 4) there was a relationship among top hits, 5) only Bacteria, Archea, and Viruses were allowed as the sources of the found hits. Using HHpred, functions were assigned to 6 more proteins. A list of these proteins was provided to Prokka as a trusted set of already annotated proteins. After the second re-annotation, only ∼25% of proteins left unclassified.

Finally, pangenome analysis of pipolins gene content was carried out as detailed above and shell-core genes present in more than 15%of pipolins were analyzed by eggNog [62] and KEGG orthology database functions with Blast Koala [63].

### Cophylogeny of pipolins and host strains

As mentioned above, the alignment of concatenated genes from the core-genome was used for the phylogeny of host strains. Phylogeny of piPolB, XerC and UDG pipolin genes was generated independently. When XerC recombinase gene appeared duplicated, so only the syntenic sequence with the UDG at the right end was included in the phylogeny reconstruction.

Phylogenetic analysis of gene sequences was carried out with Modelfinder [25] upon PRANK codon aware alignment [58].The obtained trees were then used for the comparative phylogenetic analyses with RStudio (Integrated Development for R. RStudio, Inc., Boston, MA URL http://www.rstudio.com/). Briefly, phylogenetic trees were handled and pruned when required with APE [64] and tanglegrams for visual tree comparison were generated with Phytools [65]. We used the Dendextend package [66] to calculate and represent the cophenetic distances of branches within a tree and the CCC (cophenetic correlation coefficient) between trees. Finally, we used PACo (Procrustean Approach to Cophylogeny) [39] to investigate the phylogenetic congruence between trees.

## Acknowledgments

*E. coli* 3-373-03_S1_C2 isolate was kindly provided by Dr. Ellen Silbergeld (John Hopkins University, USA). We also thank Dr. Mart Krupovic (Pasteur Institute, France) for inspiring discussions and critical reading of the manuscript. Technical assistance of María Pilar Bea-Escudero is also acknowledged.

## Funding

This research was funded by the Spanish Ministry of Science, Innovation and Universities, grant number PGC2018-093723-A-100 (AEI and FEDER, UE) to M.R.R., and by institutional grants from Fundación Ramón Areces and Banco de Santander to the Centro de Biología Molecular Severo Ochoa.

LREC laboratory was supported by projects PI16/01477 from Plan Estatal de I+D+I 2013-2016, Instituto de Salud Carlos III (ISCIII), Subdirección General de Evaluación y Fomento de la Investigación, Ministerio de Economía y Competitividad (Gobierno de España) and Fondo Europeo de Desarrollo Regional (FEDER); and ED431C2017/57 from the Consellería de Cultura, Educación e Ordenación Universitaria (Xunta de Galicia) and FEDER. S.-C. F.-S. was holder of a PhD fellowship (FPU15/02644) from the Secretaría General de Universidades, Spanish Ministerio de Educación, Cultura y Deporte.

## Author Contributions

Conceptualization (MRR), Formal analysis (SCFS, MdT, LC, MRR), Funding Acquisition (JB, MS, MRR), Investigation (SCFS, LC, MdT, MB, JMG, MRR), Resources (JB, MdT), Supervision (MdT, MS, JB, MRR), Writing – Original Draft Preparation (MRR), Writing – Review & Edit (SCFS, MdT, LC, JB, MRR).

## Supplementary Figure Legends

**Figure S1. Phylogeny of piPolB genes from analyzed pipolins**.

Nucleotide sequence of piPolB genes from new pipolins in LREC strains (cyan) and those retrieved from GenBank (orange) were aligned using Prank codon aware option and then used for maximum-likelihood phylogeny reconstruction with IQtree. Modelfinder Best-fit model was K3Pu+F+R2.

**Figure S2. Roary pangenome analysis of pipolins-harboring *E. coli* strains**.

A. Combined layout of hierarchical clustering of gene presence/absence for LREC *E. coli* genomes (left), along with color-coded markers (middle) and chromosome genetic structure (right). Representation of Roary output was rendered at Phandango website [67]. Lower panels show the accumulation of new vs. unique genes per isolate (B and D) and conserved vs. total genes per isolate (C and E) in the new (LREC, B and C) and all (LREC+GenBank, D and E) *E. coli* strains.

**Figure S3. Maximum-likelihood phylogeny of pipolins-harboring *E. coli* strains**. Codon aware core-genome alignment from Roary was used for the phylogeny reconstruction using IQtree. The best-fit model was GTR+F+R7 (according to ModelFinder). Scale bar indicates the substitution rate per site. The main features retrieved from Enterobase are indicated on the right: source, multilocus sequence type (ST) and clonal complexes (CC). Strains are colored based on the phylogenetic groups, with LREC strains highlighted in italics. Reference strain 3-373-03_S1_C2 is highlighted with a black asterisk (*).

**Figure S4. Single Nucleotide Polymorphism (SNP) trees of the pipolin-harboring strains**.

A. LREC strains’ SNP-based tree. Previously described pipolin-harboring isolate 3-373-03_S1_C2 was included as the reference genome. The SNP-tree was performed with EnteroBase [21] default parameters (min. 95% sites present) and included 132985 variant sites. Source and main clonal complexes are also indicated. B. SNP-based tree of the 58 pipolin-harboring strains available from Enterobase and the 25 LREC pipolin-harboring strains performed with EnteroBase. The tree included 170702 variant sites. The figure also includes the country of isolation, sequence types (ST) and clonal complexes (CC). Clade branches were colored by phylogenetic groups as in previous figures.

**Figure S5. Genetic structure of pipolins**.

Protein-coding genes are represented by arrows, indicating the direction of transcription, and colored as indicated in the legend. The image was generated by EasyFig software and re-annotated pipolins sorted according to the hierarchical clustering of the gene presence/absence. The greyscale on the right reflects the percent of amino acid identity between pairs of sequences. Names of pipolin-carrying strains are colored based in the phylogroups as in Figure S3.

**Figure S6. Tanglegram of pipolins and host strains**.

Tanglegram representation of maximum-likelihood comparative phylogenies of piPolB gene (same as in Figure S1) with XerC (A) and UDG (B) genes of pipolins. Association lines are colored based on the phylogenetic groups as in Figure S3.

**Figure S7. Detailed cophylogeny analysis of pipolins and host strains with PACo**. Squared Procrustes residues of each phylogenetic group were compared with the remainder interactions (A, B, C, D). The p-value of Welch t-test is indicated. Panel E shows the contribution of each pipolin-host association to the general pattern of coevolution. Each bar represents a jack-knifed estimate of a squared residual. Error bars represent the upper 95% confidence intervals from applying PACo to patristic distances. The dashed line indicates the median squared residual value that can serve as a threshold for congruent phylogenetic interactions. Bars are colored based on the phylogenetic groups of the strains.

**Table S1. Characterization of pipolin-harboring LREC strains**.

Compilation of main features determined for LREC strains. Biosample and Enterobase Uberstrain reference IDs are also indicated. D*: *Does not match with PCR data. N.D. : non-detected. * Virulence factors detected by PCR screening and in silico analysis are detailed

**Table S2. Genbank genomes carrying pipolins**.

References for genome drafts or assemblies and Biosamples as well as main features from Enterobase [21] are indicated.

*CRISPR/Cas cassettes and Integrons were surveyed as indicated in Methods.

**Integrity of detected integrons were analyzed with IntegronFinder [56] as indicated in Materials and Methods. C, complete; In0, integron lacking attC site; CALIN, integron lacking functional integrase gene.

***When available, PubMed ID of strain reporting publication

**Table S3. Functional characterization of the most common pipolin genes**.

Annotation of genes from Roary shell-genome (present in more than 15% of pipolins) is indicated. Functional groups in eggNOG and KEGG databases, as well as HHPred searches were also performed for a detailed functional characterization.

